# The ecological relevance of flagellar motility in soil bacterial communities

**DOI:** 10.1101/2024.01.22.576697

**Authors:** Josep Ramoneda, Kunkun Fan, Jane M. Lucas, Haiyan Chu, Andrew Bissett, Michael S. Strickland, Noah Fierer

## Abstract

Flagellar motility is a key bacterial trait as it allows bacteria to navigate their immediate surroundings. Not all bacteria are capable of flagellar motility, and the distribution of this trait, its ecological associations, and the life history strategies of flagellated taxa remain poorly characterized. We developed and validated a genome-based approach to infer the potential for flagellar motility across 12 bacterial phyla (26,192 genomes in total). The capacity for flagellar motility was associated with a higher prevalence of genes for carbohydrate metabolism and higher maximum potential growth rates, suggesting that flagellar motility is more prevalent in resource-rich environments due to the energetic costs associated with this trait. To test this hypothesis, we focused on soil bacterial communities, where flagellar motility is expected to be particularly important given the heterogeneous nature of the soil environment. We applied a method to infer the prevalence of flagellar motility in whole bacterial communities from metagenomic data, and quantified the prevalence of flagellar motility across 4 independent field studies that each captured putative gradients in soil carbon availability (148 metagenomes). As expected, we observed a positive relationship between the prevalence of bacterial flagellar motility and soil carbon availability in each of these datasets. Given that soil carbon availability is often correlated with other factors that could influence the prevalence of flagellar motility, we validated these observations using metagenomic data acquired from a soil incubation experiment where carbon availability was directly manipulated with glucose amendments, confirming that the prevalence of bacterial flagellar motility is consistently associated with soil carbon availability over other potential confounding factors. Flagellar motility is a fundamental phenotypic trait for bacterial adaptation to soil, defining life history strategies primarily associated with resource availability. More generally, this work highlights the value of combining genomic and metagenomic approaches to expand our understanding of microbial phenotypic traits and reveal their general environmental associations.

## Introduction

Microorganisms navigate their environment by responding to gradients in nutrients, toxins, and environmental conditions, in a process called chemotaxis [1, 2]. Flagellar motility is a widespread adaptation that allows bacteria to colonize new micro-environments by facilitating access to space and nutrients [3, 4], and enables escape from unfavorable conditions [5] and predators [6]. For example, moving towards environmental cues is an effective mechanism by which most pathogens [7, 8] and symbionts [9, 10] colonize their hosts. Despite the recognition that swimming and swarming (the two main modes of flagellar motility) are widely used to navigate microbial environments, empirical knowledge on the environmental conditions where bacterial flagellar motility can be beneficial remains rather limited, as most knowledge derives from laboratory-based studies using model organisms.

In laboratory conditions, bacteria have been widely investigated for their ability to swim towards resources [11, 12], display quorum sensing [13], or swim away from toxins [14]. Several experimental studies show that the hydration level of surfaces generally predicts how easily bacteria can colonize a given surface [15], and that flagellar motility also predicts the temporal persistence of bacterial pathogens in host microbiomes [16]. The high energetic cost of powering the flagellar machinery is tightly linked to regulatory systems that control flagellar expression depending on the spatial proximity and quality of available resources (i.e., optimal foraging based on energetic constraints; [17–19]). Notably, different flagellar systems have evolved in response to distinct environmental conditions, as exemplified by the case of the *Vibrio* genus, which use different flagellar systems depending on the spatial complexity of their surroundings [20]. This broad body of knowledge leads to the expectation that flagellar motility should display general ecological associations, but such patterns have not been comprehensively explored.

Research on bacterial flagellar motility predates modern microbiology, and laboratory-based approaches have enabled the discovery of the genes involved in flagellar assembly [21, 22]. The genes encoding for the flagellar machinery are reasonably well-known and generally conserved across a broad diversity of bacterial groups [23, 24]. Because the production of flagella requires a well-defined gene repertoire, the prediction of flagellar motility across taxa is likely feasible [25]. Yet, the proportion of bacterial taxa for which flagellar motility could theoretically be inferred from genomic information contrasts with the relatively limited number of strains for which flagellar expression has been empirically determined. A comparison of the number of strains with known motility information in bacterial phenotypic trait databases [26] to the total number of genomes contained in the Genome Taxonomy Database (GTDB r207, [27]) highlights that we have information on whether taxa are flagellated or not for only ~10% of the bacterial strains with available whole genome information. If we could infer the capacity for flagellar motility across a broad diversity of microbial taxa, we could determine the set of traits that generally characterize flagellated taxa (so-called life history strategies; [28, 29]). Previous studies have linked flagellar motility to a fast growth (copiotroph) strategy [30], and flagellar motility is expected to be associated with a life history strategy for rapid nutrient acquisition [31]. However, one of the main challenges with identifying the life history strategies of bacteria remains the quantification of phenotypic traits. Thus, developing methods to infer flagellar motility across single bacterial genomes and metagenomes can help us identify the main ecological and life history associations of this important trait.

Flagellar motility is likely common for bacteria living in many environments – including host-associated and aquatic environments [2]. However, we are particularly interested in the prevalence of flagellar motility in soil environments because soil is a heterogeneous environment where resources are patchily distributed, and access to resources is a key factor structuring soil bacterial communities [32, 33]. We expect a high degree of variability in the prevalence of motile bacteria in soil as motility requires continuous water films [34, 35], and the high energetic cost associated with flagellar motility may be disadvantageous in the resource-limited conditions often common in soil [17, 36]. Since organic carbon compounds are likely the main sources of energy for soil bacteria, soil C availability is likely a key factor determining the selective advantage of flagellar motility in soil. Indeed, several studies have found a higher prevalence of bacterial flagellar motility in soil environments that generally have higher C availability. For example, plant rhizospheres usually contain elevated levels of available C compared to adjacent bulk soil environments due to plant-derived organic carbon inputs [37], and generally harbor a higher prevalence of flagellar genes [38, 39]. In arid environments, several studies have detected a negative relationship between the prevalence of flagellar motility genes and aridity [40, 41], which could be due to both lower C availability or to lower moisture. Given the spatial heterogeneity of soil, and the fitness advantage theoretically gained from flagellar motility in conditions where energy-rich resources are patchily distributed [18], we hypothesize that bacterial flagellar motility should exhibit a general positive relationship with soil C availability.

We had three objectives with this study. First, we wanted to build genome and metagenome-based models to accurately infer the potential for flagellar motility across bacterial taxa and whole bacterial communities. Second, we sought to identify the general life history strategies associated with flagellar motility in bacteria. Third, we aimed to determine the prevalence of flagellar motility in soil bacterial communities and to test the hypothesis that flagellar motility is more prevalent in soils with higher C availability. To this end, we estimated the potential for flagellar motility across 26,192 bacterial taxa with available genomic information based on a machine learning model trained on empirical information for this trait, and explored whether flagellar motility is associated with broader life history strategies. We then applied a method to estimate flagellar motility as a community-aggregated trait directly from metagenomes. We used this method to investigate the prevalence of flagellar motility across four independent sample sets that we would expect to capture gradients in soil C availability, and confirmed our findings using metagenomes from a soil incubation experiment where C availability was experimentally manipulated via glucose amendments.

## Results and Discussion

### Development of a genomic model to predict flagellar motility in bacteria

Since the genes involved in flagellum assembly are well-described and conserved across bacterial groups [23], we were able to use information on the presence/absence of flagellar genes to predict the capacity for flagellar motility from genomic information alone. We used genomic data for 1225 bacterial strains known to be motile or non-motile using strain description information compiled in [26] as training data for a boosted regression machine learning model to predict the capacity for flagellar motility in bacteria (388 unique strains with known flagellar motility and 837 unique strains with no flagellar motility; Supplementary Data 1). We note that being non-flagellated does not mean taxa are non-motile. For example, within the Bacteroidota, which had only 2 flagellated members in the training data, the majority of aquatic and terrestrial members display gliding motility [42, 43]. Of the initial set of 35 genes we identified from the literature as being associated with flagellum assembly (Supplementary Data 2), we found that 14 out of these 35 genes were either not frequently found in the genomes of taxa with experimentally validated flagellar motility, or occurred in >50% of the genomes of non-flagellated taxa. As these 14 genes were not useful for predictive purposes, the final model was based on the presence/absence of 21 genes that were sufficiently prevalent across bacterial genomes and less frequently found in non-motile taxa (Supplementary Data 2). These genes encode different structural parts of the flagellar apparatus, including the basal body (*FlaE*, *FliL*, *Flg_bbr_C*, *Flg_bb_rod*), the flagellar rotor (*FliG_C*), the flagellar hook (*FlgD*, *Flg_hook*, *FliD_C*, *FliE*), or the M-ring (*YscJ_FliF_C*), as well as multiple proteins for protein export and the flagellins required for flagellar assembly (Supplementary Data 2). We verified that the presence/absence of this set of genes could effectively distinguish taxa with flagellar motility from non-flagellated taxa using Principal Components Analysis (PCA) (Supplementary Figure 1A).

Our model inferred that taxa were able to display flagellar motility correctly in all taxa with experimentally verified flagellar motility, and inferred that taxa were non-flagellated correctly in 94.5% of the cases (Supplementary Figure 1B). We verified that many of the genomes that the model incorrectly predicted as having flagellar motility belonged to strains whose genomes contain the majority of flagellar motility genes and have sister taxa that do display flagellar motility (Supplementary Data 3). We also recognize that a number of strains might express flagella under certain environmental conditions that would not be captured with the specific *in vitro* conditions used for strain isolation and phenotyping. Unsurprisingly, the phylum Proteobacteria was overrepresented in the phenotypic trait database (Supplementary Figure 2), but our predictions of flagellar motility for this phylum were not necessarily more accurate than predictions for other phyla (Supplementary Figure 3), as it contained numerous taxa considered to be non-motile with flagellated sister taxa (Supplementary Data 3). We recognize that our dataset is over-represented by taxa (particularly those within the Proteobacteria, Actinobacteria, and Firmicutes) that are readily cultivated *in vitro* as those are the only taxa for which phenotypic information on flagellar motility is available. However, given that the genes associated with flagellar motility are generally well-conserved across a broad diversity of bacteria [23], and given that our model was robust across multiple phyla (Supplementary Figure 3), we expect that our genome-based model can also effectively predict flagellar motility for taxa with no available phenotypic information on motility, including taxa not included in our test set.

### Prevalence of flagellar motility across a broad diversity of bacteria

We next used our validated genome-based model (based on the presence/absence of 21 genes) to determine how the potential for flagellar motility is distributed across a broad diversity of bacteria, including a wide range of taxa for which no phenotypic information on motility is currently available. We did so to assess the degree to which flagellar motility is predictable based on taxonomic or phylogenetic information, and to investigate the genomic attributes that are generally associated with flagellar motility. We predicted the capacity for flagellar motility for 26,192 bacterial genomes spanning 12 major phyla (covering all high-quality genomes in GTDB r207 [27], belonging to the main bacterial phyla; see Methods). The predicted prevalence of flagellar motility was highly variable among phyla, ranging from the phylum Spirochaetota, which had the highest proportion of flagellated taxa (93.2%) to the Deinococcota and Mycoplasmatota which our model suggests do not have any flagellated members (Figure 1A). Among the phyla with the largest number of genomes, we found that the Proteobacteria are predominantly flagellated (78.3%), with lower proportions for the Firmicutes (54.6%), and very low proportions of flagellated taxa in the phyla Actinobacteriota (15.9%) and Bacteroidota (0.7%) (Figure 1A).

**Figure 1.**
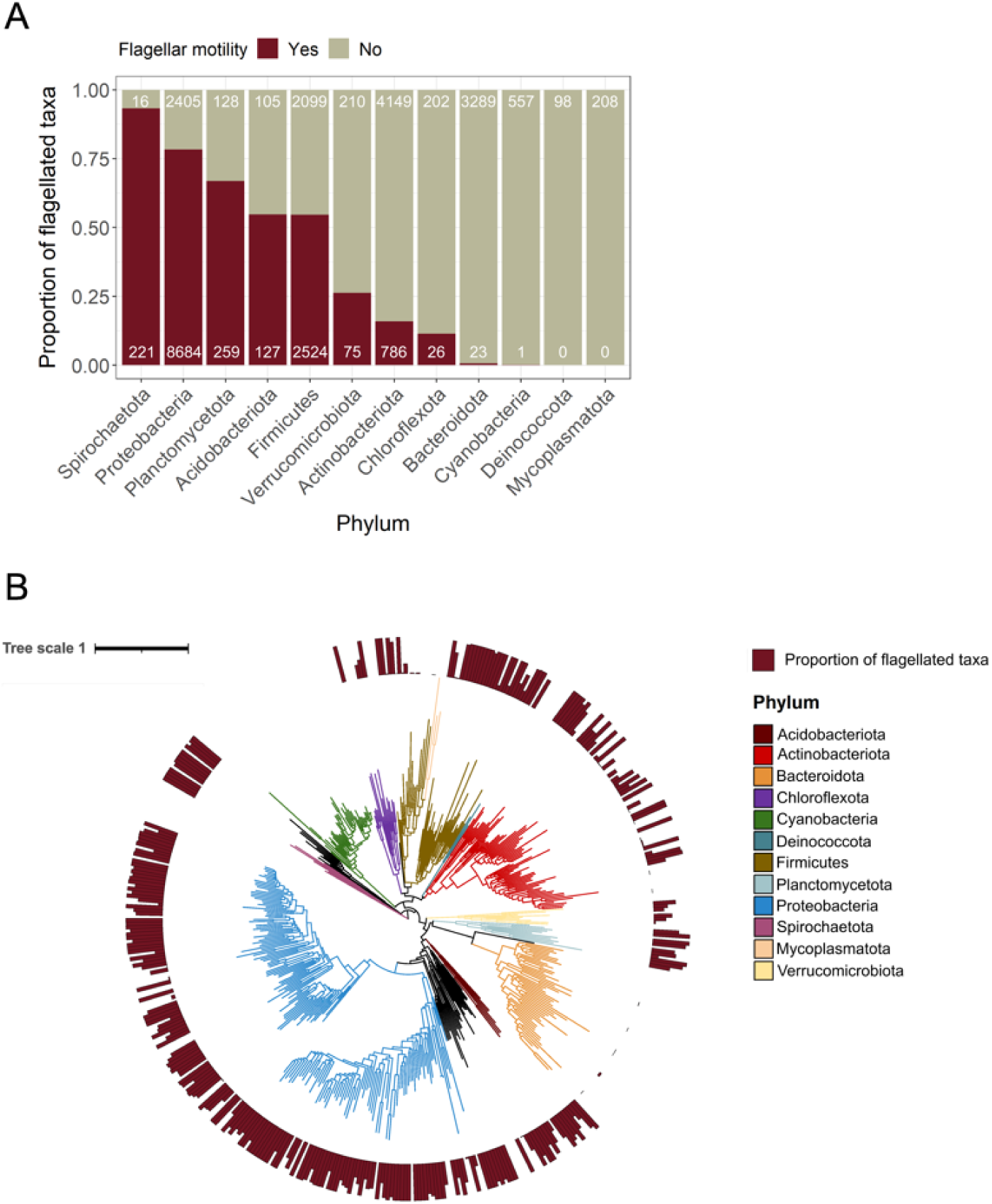
Taxonomic and phylogenetic distribution of flagellar motility in bacteria. A. Prevalence of flagellar motility in bacterial taxa from the 12 phyla best-represented phyla in a curated database of reference genomes (N = 26,192 genomes). B. Phylogenetic distribution of flagellar motility across the 12 bacterial phyla. To construct the tree, we randomly selected a single genome representative of each family found in each phylum, and predicted the capacity for flagellar motility in these genomes. Higher bars indicate a greater proportion of genomes within that family that are inferred to have the capacity for flagellar motility (based on our genome-based model, see Methods). The tree was constructed from the Genome Taxonomy Database phylogeny (GTDB r207; [27]).

The majority of bacterial phyla contain numerous families with both flagellated and non-flagellated members (Supplementary Figure 4). This means that family-level taxonomic information alone cannot necessarily provide robust inferences of flagellar motility, stressing the need for alternative approaches to evaluate the prevalence of this trait across microbial communities. However, at the genus level, most taxa are either flagellated or non-flagellated, indicating that this trait is typically conserved at this level of taxonomic resolution (Supplementary Figure 5).

Consistent with our taxonomic analyses, the phylogenetic analyses also highlight that the prevalence of flagellar motility is highly variable at broad taxonomic levels, which was reflected in a weak phylogenetic signal (phylogenetic *D* = −0.077, P < 0.001; Figure 1B). Higher-resolution phylogenetic and taxonomic information can often be useful for inferring flagellar motility, particularly for those groups that are well-characterized (i.e., where information on flagellar motility, or lack thereof, is available for closely related taxa). However, phenotypic information is often unavailable for the broad diversity of taxa found in environmental samples, highlighting the utility of the genome-based predictive approach described here that makes it feasible to leverage the rapidly expanding databases of bacterial genomes to comprehensively investigate the prevalence of this trait in microbial communities.

### Is there a general life history strategy associated with flagellar motility in bacteria?

We expect that bacteria with the capacity for flagellar motility should have distinct ecologies from non-flagellated taxa. In particular, we expect that flagellated taxa should be capable of more rapid growth and a greater capacity for carbohydrate degradation than non-flagellated taxa (i.e. a ‘resource-acquisition’ life history strategy; [31]). By analyzing the 26,192 genomes for which we had inferred the capacity for flagellar motility, we were able to identify genomic attributes that were consistently associated with flagellar motility, conducting these analyses separately for each of the 6 phyla which had sufficient representation of both flagellated and non-flagellated taxa (Figure 2A; see Methods). Besides the expected overrepresentation of genes for motility and extracellular structures (Figure 2A), the two gene categories that were consistently over-represented in taxa with the capacity for flagellar motility were signal transduction mechanisms (linked to chemotaxis) and carbohydrate transport and metabolism (Figure 2A). The latter observation is consistent with our general expectation that flagellar motility should be associated with a ‘resource-acquisition’ life history strategy (sensu [31]). However, we note that this pattern was only evident in 4 of the 6 phyla examined (Figure 2A).

**Figure 2.**
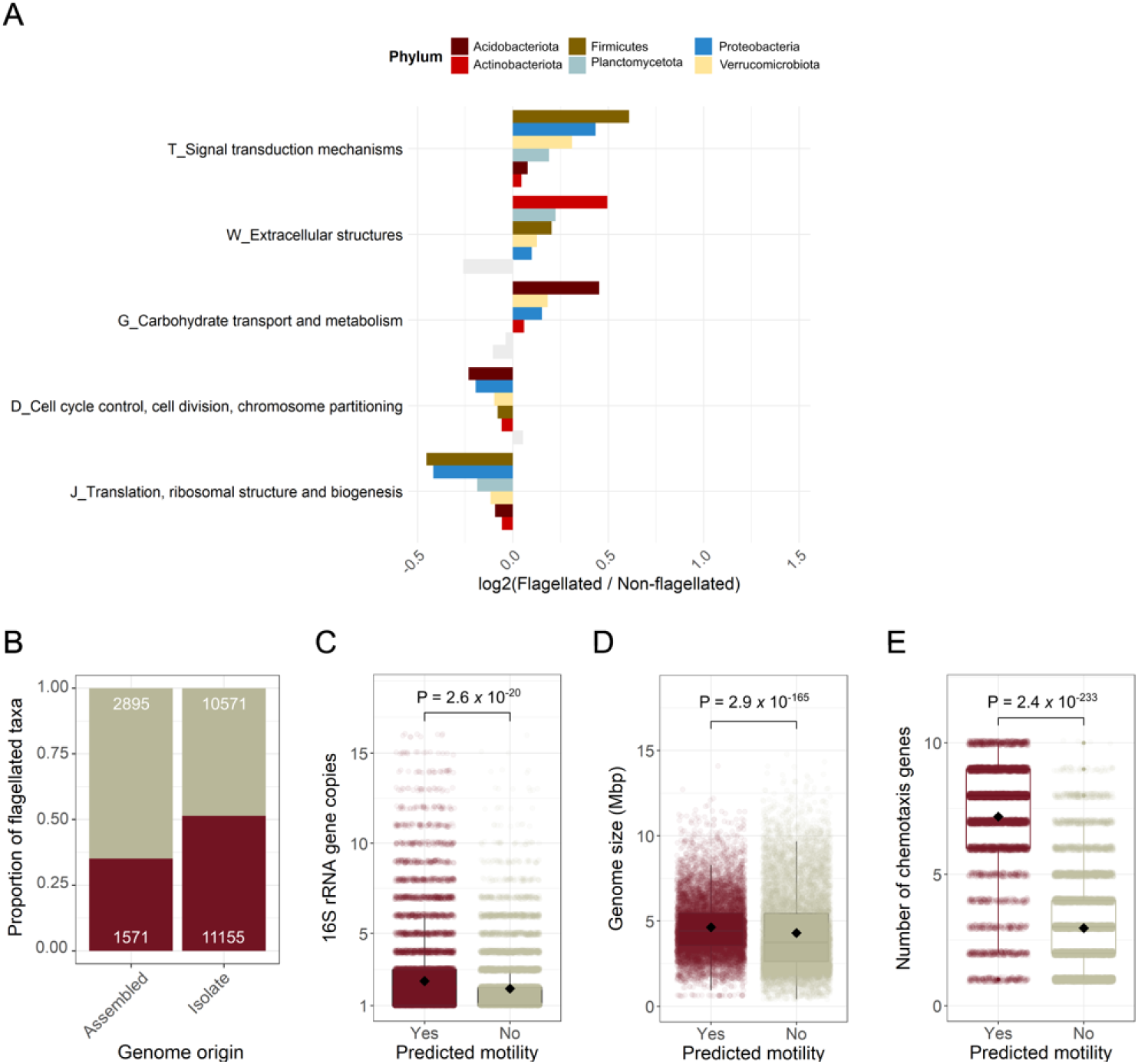
Genomic attributes associated with bacteria inferred to have the capacity for flagellar motility. A. Functional gene categories that are consistently overrepresented in genomes from taxa predicted to be flagellated or non-flagellated across the 6 most dominant phyla that contain >15% flagellated taxa. Functional categories were defined as Clusters of Orthologous Genes (COGs). We indicated those gene categories that were not statistically different with grey shading based on Mann-Whitney U tests (P > 0.01). B. Prevalence of flagellar motility in genomes derived from environmental metagenomes (MAGs) or single cells (SAGs) (‘Assembled’), and in genomes obtained from bacterial isolates (‘Isolate’). Numbers on the upper and lower ends of the plot indicate the number of genomes predicted to be non-flagellated and flagellated, respectively. C. Number of 16S rRNA gene copies in genomes of taxa predicted to be flagellated (N= 12,726) versus non-flagellated (N = 13,236). D. Genome size of taxa predicted to be flagellated and non-flagellated. E. Number of genes involved in chemotaxis identified in the genomes of taxa that are flagellated and non-flagellated. In panel A, N_Acidobacteriota_ = 232; N_Actinobacteriota_ = 4935; N_Firmicutes_ = 4623; N_Planctomycetota_ = 387; N_Proteobacteria_ = 11,089; N_Verrucomicrobiota_ = 285. In panels C and E the P-value was obtained from Mann-Whitney U tests due to non-normality of the data. In panel D, the P-value was obtained from a Welch two-sample t-test, (P < 0.05); N = 26,192 genomes.

We also found that 51.3% of genomes obtained from cultured isolates were predicted to be flagellated, compared to only 35.2% of genomes of assembled origin (metagenome-assembled and single cell-assembled genomes, MAGs and SAGs, respectively; Figure 2B). As culture collections are generally biased towards faster growing bacterial taxa with adaptations for rapid substrate uptake [44], these results provide additional support for the hypothesis that flagellar motility is often indicative of a ‘resource acquisition’ life history strategy [31].

To complement these analyses, we also determined the total number of 16S rRNA gene copies per genome as a proxy for maximum potential growth rate in bacteria [45]. The number of 16S rRNA gene copies was significantly higher in genomes of taxa inferred to have the capacity for flagellar motility (Mann-Whitney U P < 0.001; Figure 2C), but this pattern was only significant in two phyla (Firmicutes and the Proteobacteria, Supplementary Figure 6A). Genome size was also significantly larger in taxa predicted to display flagellar motility (Mann-Whitney U P < 0.001; Figure 2D), a pattern that was consistent across all phyla except for the Actinobacteriota (Supplementary Figure 6B), and agrees with previous work [29]. We additionally verified that flagellated taxa harboured a significantly higher number of genes for chemotaxis than taxa predicted to be non-flagellated (Figure 2E), as we would expect (Figure 2A; [4]).

Together, our genomic analyses suggest that bacteria with flagellar motility tend to be capable of more rapid growth and the rapid acquisition of organic C substrates, but this pattern is variable across phyla. Consistent with our findings, a recent global classification of life history strategies in bacteria found flagellar motility to be associated with elevated genomic capacity for carbohydrate metabolism, higher 16S rRNA gene copy numbers, and larger genomes [29]. Recent studies focusing on soil bacterial communities have had similar findings [38, 46], and in aquatic environments flagellar motility is considered a signature of copiotrophic lifestyles [30]. Overall, our findings suggest that flagellar motility is often part of a general life history strategy for rapid organic carbon metabolism and high maximum potential growth [31], recognizing that these analyses are based on a biased subset of bacterial diversity [44] given that most of the genomes included in this analysis (83%) were derived from cultivated isolates.

### Application of a metagenome-based approach to quantify the prevalence of flagellar motility in bacterial communities

We next extended our genome-based method so it could be used to infer the prevalence of flagellar motility in whole communities. As the prevalence of flagellar motility is difficult to reliably infer from taxonomic information alone (see above), and because neither genomic data nor phenotypic information is available for many environmental bacteria, we used a metagenome-based method to quantify flagellar motility as a community-aggregated trait [47]. This method is based on calculating the ratio between the 21 genes identified as being indicative of flagellar motility and single-copy marker genes detected per metagenome (see Methods and overview provided in Figure 3A). We first validated this metagenomic approach using simulated metagenomic data (see Methods). The simulated data were derived by mixing different proportions of genomes from taxa with experimentally-verified flagellar motility capabilities, creating a gradient of metagenomes containing between 0% and 100% flagellated taxa (Figure 3B). This allowed us to obtain a linear equation to predict the prevalence of flagellated bacteria in any given metagenome based on the summed abundances of the 21 genes indicative of flagellar motility (as determined from the genomic analyses above) to the summed abundances of single-copy genes shared across nearly all bacteria (using a similar approach to [48]; Figure 3A). With these simulated metagenomes, the ratio between the median gene length-corrected reads per kilobase assigned to flagellar and single-copy marker genes was strongly correlated with the proportion of flagellated taxa in bacterial communities assembled *in silico* (Pearson’s correlation r = 0.99, P < 0.0001; Figure 3B). We further validated the approach with metagenomic data obtained by sequencing a DNA mixture from the commercial ZymoBIOMICS microbial community standard, which contains known amounts of genomic DNA from different bacterial taxa whose flagellar motility capabilities are known *a priori* (see Methods). We found that this method accurately inferred the proportion of taxa that were flagellated based on metagenomic information alone (estimated proportion of flagellated taxa = 52.0%, expected proportion of flagellated taxa = 48.2%; Figure 3B). We also verified that our estimates using only the forward reads did not differ from those using the reverse or merged reads (Figure 3B). Together, these results highlight that we can accurately infer the community-level prevalence of bacterial flagellar motility in any metagenome of interest simply by calculating the ratio between the sum of the 21 flagellar genes and the sum of single-copy bacterial marker genes.

**Figure 3.**
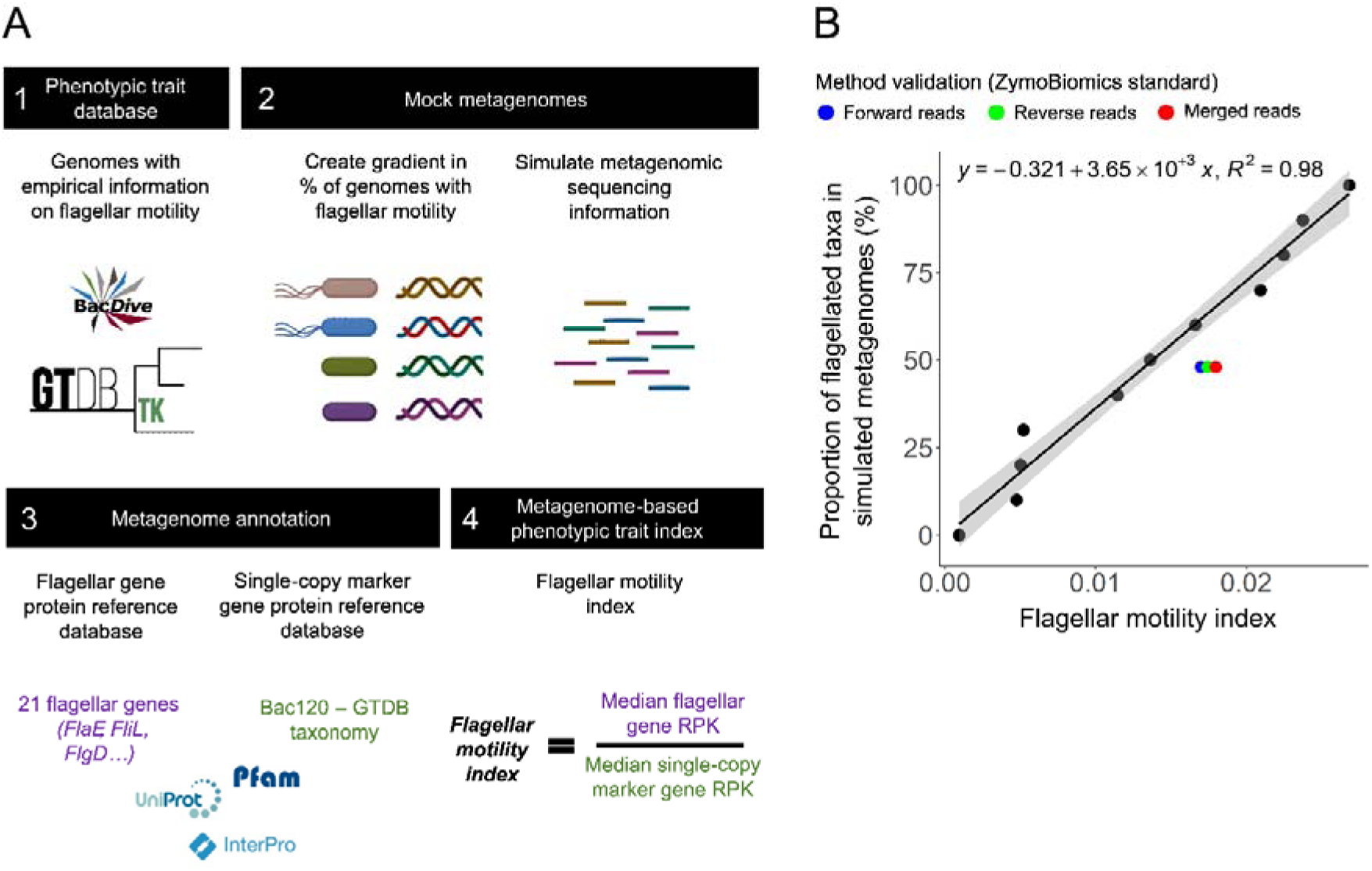
Developing a metagenome-based approach to quantify the prevalence of flagellar motility in bacterial communities. A. Method overview. We first collected whole-genome data for bacterial taxa directly observed to have flagellar motility *in vitro* (1). We then make combinations of genomes with and without the capacity for flagellar motility to create a gradient of the prevalence of flagellar motility in ‘mock’ metagenomes (2). These ‘mock’ metagenomes are created by simulating shotgun metagenomic reads from the whole genomes (see Methods). We annotate the metagenomes to identify the 21 flagellar genes determined from the genomic analyses to be indicative of flagellar motility along with a set of 120 single-copy marker genes that are found in nearly all bacteria (see Methods) (3). Finally, we calculate the gene length-corrected reads-per-kilobase (RPK) of each of these gene sets and calculate a ‘flagellar motility index’ using the ratio between these indices (4). B. Linear relationship between the ‘flagellar motility index’ calculated as shown in panel A (4) and the proportion of genomes of taxa with flagellar motility in simulated metagenomes (panel A, 2) (N = 14). The y-axis shows a gradient of bacterial metagenomes created by combining different proportions of genomes from bacteria known to be flagellated or non-flagellated spanning the phyla Proteobacteria, Actinobacteria, and Firmicutes. The linear equation resulting from this association can be used to quantify the prevalence of flagellar motility in any bacterial metagenome. Colored dots indicate the known proportion of flagellated taxa in the metagenome of the ZymoBiomics microbial community standard.

### Prevalence of bacterial flagellar motility across gradients in soil carbon availability

We used our metagenome-based approach to further test our hypothesis that flagellar motility is most likely to be associated with taxa adapted for fast resource acquisition under resource-rich conditions (Figure 2A-C). If this hypothesis is valid, we would expect the community-wide prevalence of bacterial flagellar motility to be higher in soils with greater amounts of available organic C. Since it is challenging to directly quantify the amount of C in soil that is available to fuel microbial activities, we selected 4 independent metagenomic datasets that we expect to effectively capture gradients in soil C availability, and the results from the analyses of these datasets are described below.

Soil C availability is expected to decrease with soil depth [49, 50]. Across the 9 soil depth profiles analyzed [51], we consistently observed a higher prevalence of flagellar motility in the surface (top 20cm) compared to deeper soil horizons (20-90cm, linear mixed effects model, Estimate_Surface_ = 11.88 ± 1.30 (mean ± SD), Estimate_Subsurface_ = 8.64 ± 1.34, P = 0.005, N = 66; Figure 4A; Supplementary Figure 7). We recognize that soil C availability is not the only factor that is likely to change appreciably with soil depth. For example, soil water and nutrient availability can also vary with depth [51], so we cannot conclude that soil C availability is the only factor responsible for the elevated prevalence of flagellar motility in surface soil communities.

**Figure 4.**
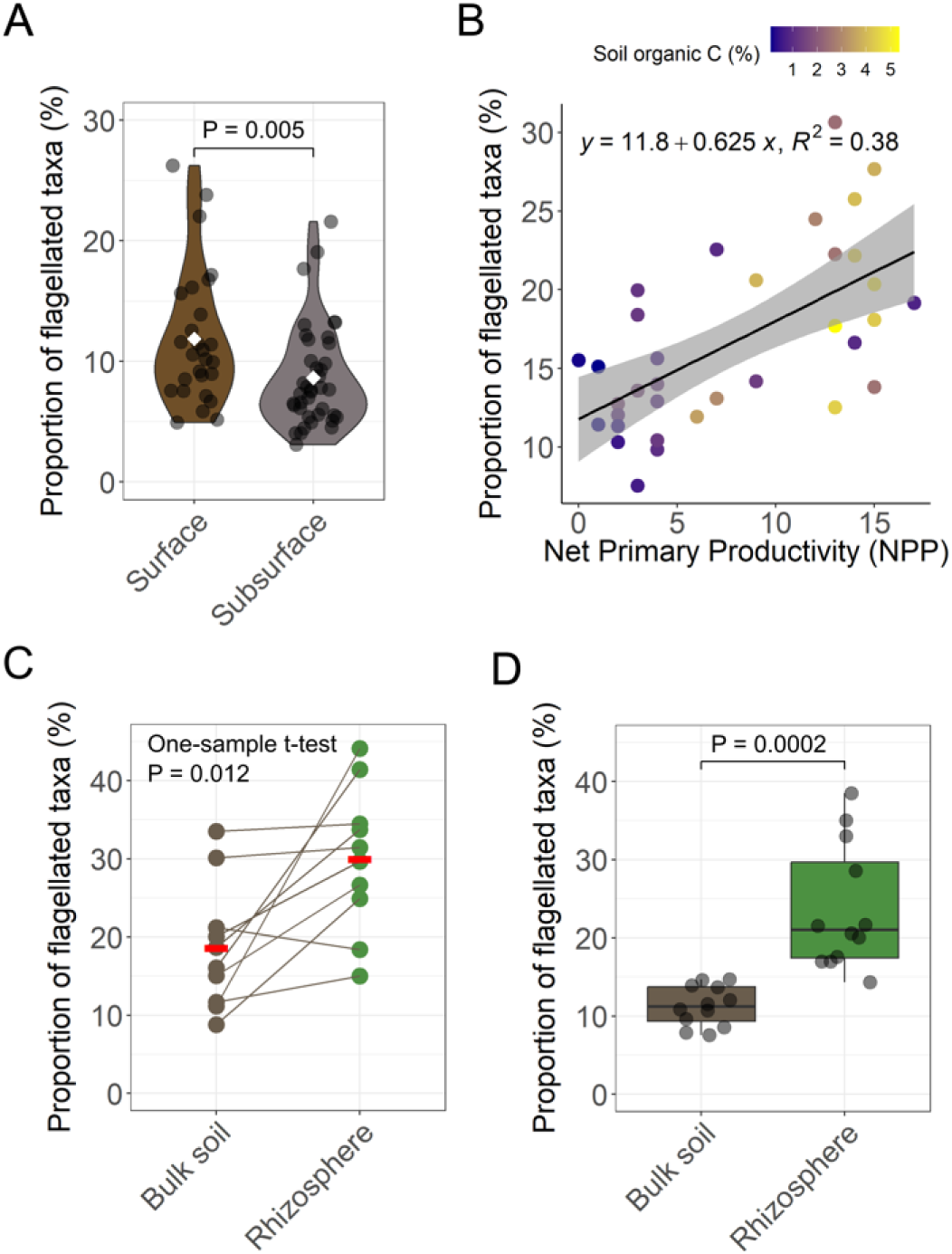
Prevalence of flagellar motility in bacterial communities spanning putative gradients in soil carbon (C) availability. A. Estimated prevalence of flagellar motility in bacterial communities found across soil profiles (Surface, 0-20cm; Subsurface, 20-90cm; N = 66). These soil profiles were sampled from sites that covered diverse climatic regions across the USA [51]. Group means are shown as white diamonds, and the P-value was obtained from a linear model with site coded as a random factor. B. Relationship between the estimated prevalence of flagellar motility and net primary productivity (NPP) across Australia (N = 38; [81]). The shaded area depicts the standard error around the mean. C. Comparison of the prevalence of flagellar motility in bulk soils and rhizospheres of citrus trees found at 10 sites across the globe (N = 20; [55]). Since each site contained a single bulk soil and a single rhizosphere sample, we indicate which samples come from the same site using connecting lines. To obtain the P-value, we calculated the difference between the prevalence of flagellar motility in the rhizosphere and bulk soil at each site, and then made a comparison against zero using a one-sample t-test. Means are shown as horizontal red lines. D. Comparison of the prevalence of flagellar motility in bulk soils and rhizospheres of wheat plants from a controlled pot experiment (N = 24; [56]). The P-value was obtained using a Welch two-sample t-test. Statistical significance is set at P < 0.05.

To further test our hypothesis, we analyzed 38 surface soils collected from across Australia. For this sample set, we assume that net primary productivity (NPP) is a reasonable proxy for soil C availability, as higher NPP leads to increased plant-derived organic matter inputs to soil [52]. We found that across these varied soils, the prevalence of flagellar motility in bacterial communities was strongly correlated with NPP (Pearson’s r = 0.619, P < 0.001; Figure 4B). As with the ‘soil depth’ analyses, these findings also support our hypothesis that flagellar motility is more prevalent in soils with higher C availability. However, as other factors likely co-vary with NPP (including mean annual precipitation), these findings on their own are not sufficient to confidently support a general association between flagellar motility and soil C availability.

As both the ‘soil depth’ and the ‘Australian surface soil’ datasets indicate an association between inferred soil C availability and flagellar motility, we then sought to determine the prevalence of flagellar motility in bacteria from rhizospheres and associated bulk soils. While many factors differ between rhizosphere and bulk soils, we would expect that soil C availability is one of the more prominent factors differing between these two soil habitats. The rhizosphere receives abundant inputs of available plant-derived C via root exudation [53], and rhizosphere soils generally support higher microbial respiration rates than adjacent bulk soils [37, 54]. We analyzed two independent metagenomic datasets that compared bacterial communities in rhizospheres and adjacent bulk soils. One dataset contained paired rhizosphere and bulk soil samples across the globe from diverse citrus species ([55], N = 20), and the other dataset contained samples from a controlled pot experiment with wheat plants ([56], N = 24). We found that in both datasets, rhizosphere bacterial communities consistently had a higher prevalence of flagellar motility compared to their adjacent bulk soils (Figures 4C,D). Across citrus species, the prevalence of flagellar motility was on average a 11.5% higher in rhizospheres than in bulk soils (one-sample t-test P = 0.012; Figure 4C; Supplementary Figure 8), and was higher in the rhizosphere than in the paired bulk soil in 9 out of the 10 sites analyzed. In wheat plants, we also found that rhizosphere bacterial communities contained a higher prevalence of flagellar motility (Estimate_Rhizosphere_ = 23.7 ± 2.6) than bulk soils (Estimate_Bulk soil_ = 11.3 ± 8.0; Welch two-sample t-test P = 0.0002; Figure 4D). While we recognize that other factors could contribute to the elevated prevalence of flagellar motility in rhizosphere communities, these results provide further support for our hypothesis that flagellar motility is favored under conditions of higher soil C availability, as also indicated by the analyses of the ‘soil depth’ and the ‘Australian surface soil’ datasets.

### Experimental verification that bacterial flagellar motility is associated with soil carbon availability

To more conclusively test whether soil C availability is associated with the prevalence of bacterial flagellar motility, we generated new metagenomic data from a 117-day soil incubation experiment where C availability was directly manipulated via regular glucose amendments (see Methods; [57]). This experiment was performed in the absence of a growing plant and under uniform moisture conditions, thus minimizing the impact of these potential confounding factors. The prevalence of flagellar motility in the bacterial communities amended with glucose (15.82 ± 0.89%) was higher than in the soils that did not receive glucose (13.19 ± 1.56%) (Welch two-sample t-test P = 0.017; Figure 5A). This pattern is in line with the results from the field studies (Figure 4) and supports our central hypothesis that the prevalence of flagellar motility is positively associated with soil C availability. The rather small size of these effects is likely due to the fact that relatively few bacterial taxa responded to the glucose addition. While glucose addition shifted the overall community composition (Figure 5B), only 28 bacterial taxa (ASVs) out of the total 1203 ASVs detected were significantly more abundant in the glucose-amended soils. These taxa that significantly responded to glucose addition belonged to 7 different bacterial phyla (Supplementary Figure 9).

**Figure 5.**
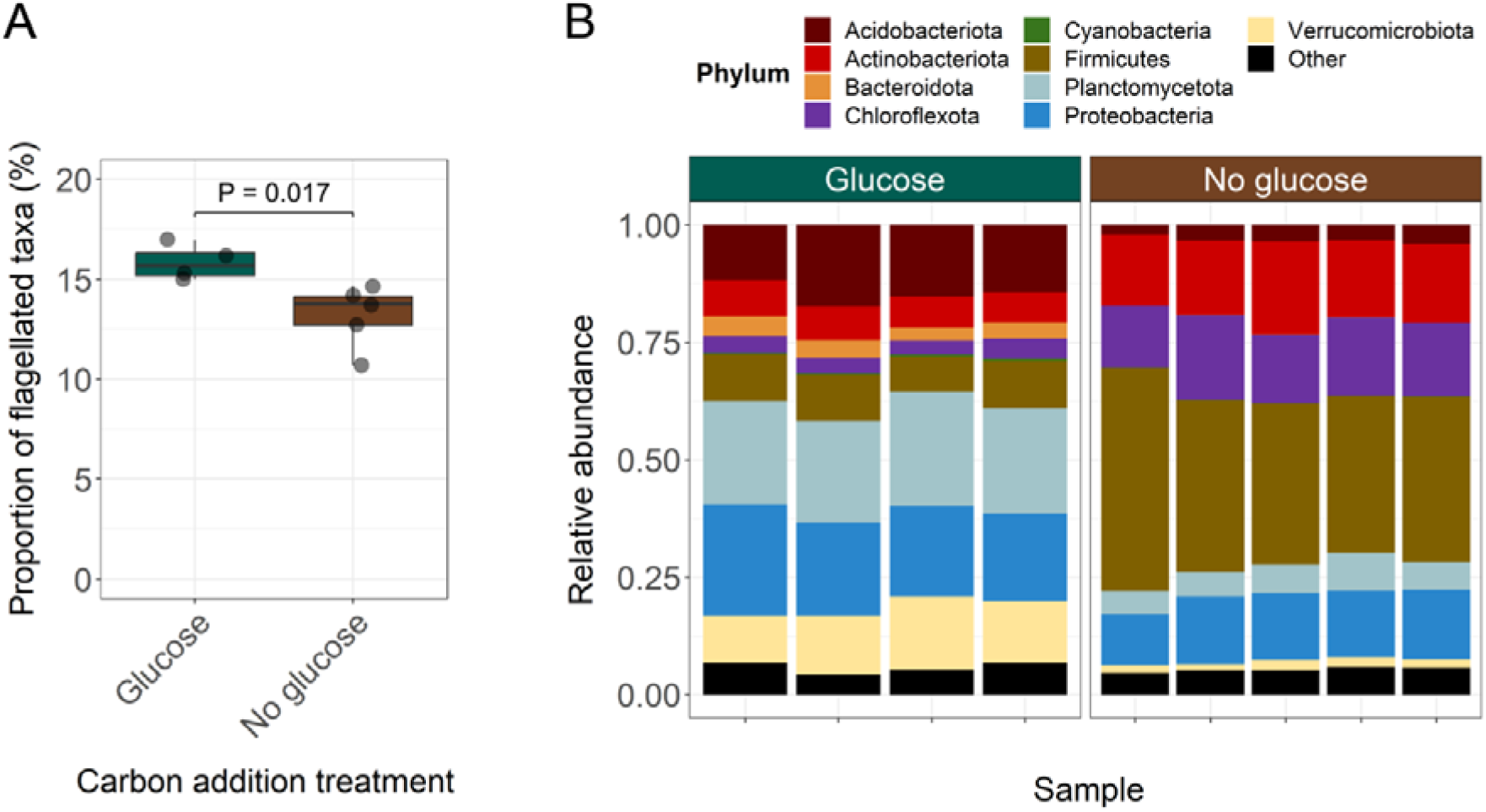
Prevalence of flagellar motility in bacterial communities from a 117-d soil incubation experiment where soil carbon (C) availability was directly manipulated via addition of glucose. A. Estimated prevalence of flagellar motility in bacterial communities found in soils amended (N = 4) and not amended (N = 5) with glucose as a way to directly manipulate soil C availability [57]. The P-value was obtained using a Welch two-sample t-test with significance at P < 0.05. B. Taxonomic composition of bacterial communities from soils amended or non-amended with glucose over 117 days of incubation. The taxonomic composition of the bacterial communities was determined via amplicon sequencing of the 16S rRNA gene (see Methods).

### Conclusions

We have shown that flagellar motility is a key trait linking C dynamics and microbial communities in soil. Consistent with expectations [31], our genomic analyses reveal that flagellated taxa tend to be associated with a ‘resource-acquisition’ life history strategy. This observation was supported by our metagenomic analyses which revealed a positive relationship between the prevalence of flagellar motility in bacterial communities and soil C availability across multiple, independent datasets. This relationship between flagellar motility and soil C availability can be explained based on fundamental energetic constraints, which make flagellar motility a beneficial trait in environments where C availability is elevated, particularly in spatially structured environments like soil where available C can be patchily distributed [35].

The methods to predict microbial traits from genomic information presented here are particularly relevant for traits that are difficult to quantify *in situ* or for those that require isolation and culturing [58, 59]. Our metagenome-based approach to infer the proportion of a microbial community harboring any given phenotypic trait would be very useful for this purpose (Figure 3A). This method can also be applied to investigate processes where flagellar motility is expected to play an important role, such as microbial colonization and persistence in host-associated microbiomes [60, 61]. In efforts to improve microbiome management, a better quantification of the prevalence of flagellar motility in these systems could help identify microbiomes that are likely to be more persistent in the host or more likely to deliver beneficial functions [62]. These methods could also be used to explore the prevalence of motility and its associated traits across gradients in C availability in other environments of interest, such as freshwater systems. Overall, genome-based predictive approaches offer opportunities for expanding our trait-based understanding of microbial communities beyond cultivated taxa, and help us understand microbial community patterns across environmental gradients.

## Materials and Methods

### Genome selection and annotation

We compiled genomic data from ~62,000 unique bacterial taxa (‘species clusters’) available in the Genome Taxonomy Database (GTDB) (release 207; [27]). We restricted our analyses to bacterial phyla with more than 100 representative genomes available in GTDB and only included genomes estimated to be >95% complete based on CheckM (v1.1.6) [63]. We also removed all genomes that lacked a 16S rRNA gene, as well as those with signals of chimerism based on GUNC (Genome Unclutterer; [64]), yielding 26,192 genomes in total belonging to 12 different phyla.

The coding sequences of the 26,192 genomes were identified using Prodigal (v2.6.3; [65]). We then aligned the predicted coding sequences for each genome to the Pfam database (v35.0; [66]) using HMMER (v3; [67]) to obtain information on all potential domains and genes present in those genomes. All matches with a bit score lower than 10 were discarded. We then binarized all copy numbers of genes and domains in each genome to presence/absence for further analyses. We selected a set of 21 genes out of a total of 35 genes involved in flagellar assembly in Pfam based on their prevalence among strains with empirical information on flagellar motility (Supplementary Data 2). Specifically, this subset of genes was chosen based on the following criteria: 1) genes were present in >80% of taxa with experimentally demonstrated flagellar motility, and 2) genes were not present in >50% of taxa classified as non-motile based on available phenotypic information (see below). This step was necessary as many non-motile taxa conserve genes for flagellar motility (Supplementary Data 3), and some of the flagellar genes are not well represented in Pfam. We used the information on the presence/absence of these 21 genes across genomes to build a predictive model of flagellar motility in bacteria (Supplementary Data 2).

### Genome-based prediction of flagellar motility in bacteria

We compiled all information on whether bacterial taxa displayed flagellar motility or not from the bacterial phenotypic trait database compiled in [26]. This database contains information on motility traits for 13,481 unique bacterial strains [26]. We first selected only the subsets categorized as having flagellar motility or as being non-motile (8191 unique strains). To obtain representative genomes for these strains, we matched the National Center for Biotechnology Information (NCBI) taxon id of each of these strains to their corresponding genome accession in GTDB. To ensure maximal reliability of the genomic information used for model training, we only kept those genomes that were 100% complete, and applied the same quality filters mentioned above. This led to a final subset of 1225 high quality genomes (388 categorized as having flagellar motility, 837 non-motile) that we used for model training (Supplementary Data 1). We note that these 1225 genomes included taxa from 18 unique phyla, with the proportions of motile taxa per phylum ranging from 0-100% (Supplementary Figure 2).

Since some of the genes involved in flagellar assembly are often present in several non-motile taxa ([68]; Supplementary Data 3), we were not able to use standard statistical approaches to build a predictive model of flagellar motility based on the presence/absence of 21 flagellar genes. We thus used gradient boosted regression decision trees that could accommodate the complexity of having 21 predictive features using the xgboost package in R (v1.7.5; [69]). To this end, we first built a training and a test set (70:30, randomly selected) of the matrix containing the presence/absence of the flagellar genes for each of the representative bacterial genomes with experimental information on flagellar motility using the *xgb.DMatrix* function of xgboost. We then applied Bayesian hyperparameter optimization to select the best parameters for the regressor model using the *bayesOpt* function of the ParBayesionOptimization R package (v1.2.6; [70]), specifying the objective function as a binary logistic regression. We ran k-fold cross-validation using the *xgb.cv* function in xgboost to identify the optimal number of iterations of model improvement for the final model training function. We built the boosted regression model using the optimized parameters and iterations calculated above using the *xgboost* function of package xgboost. We used the function *xgb.importance* from xgboost to compute the predictive importance of the different genes in the final model, which identified 14 flagellar genes that were most useful for predicting flagellar motility (Supplementary Data 4), even though the full set of 21 genes was needed for accurate prediction (see Results and Discussion for details on the performance of the final selected model). We evaluated model performance using the accuracy index.

### Phylogenetic analysis

To investigate the phylogenetic distribution of flagellar motility in bacteria, we first randomly selected a single genome from each family within the 12 predominant phyla investigated (485 genomes in total). Since we had already predicted the potential for flagellar motility across GTDB genomes, we simply subsetted the tree provided by GTDB with the selected genomes, which can be found in https://data.gtdb.ecogenomic.org/releases/release207/207.0/bac120_r207.tree. This tree is based on the alignment of 120 single-copy marker genes and is therefore more robust than a conventional maximum likelihood tree based on the alignment of full 16S rRNA gene fragments. We visualized and edited the trees using iTOL (v5; [71]). We tested whether flagellar motility had a phylogenetic signal by calculating the phylogenetic *D* index for binary traits [72], where values (positive or negative) closer to 0 indicate phylogenetic conservatism, and values closer to 1 indicate a random phylogenetic pattern. This phylogenetic analysis was conducted using the R package ape (v5.7-1; [73]). We additionally explored the degree of conservatism of flagellar motility across different levels of phylogenetic resolution by measuring the standard deviation (SD) of the flagellar motility status (flagellated, 1; non-flagellated, 0) across taxa from different taxonomic ranks (phyla, classes, orders, families, and genera). For this analysis, we only included those taxa that were represented by more than one genome.

### Analysis of bacterial life history strategies associated with flagellar motility

We investigated associations between flagellar motility and broad functional gene categories by testing the prevalence of Clusters of Orthologous Genes (COGs) in the genomes of taxa predicted to be flagellated or non-flagellated [74]. We excluded the phyla Bacteroidota, Chloroflexota, Cyanobacteria, Spirochaetota, and Mycoplasmatota from this analysis as these phyla had either too high (>90%) or too low (<15%) proportions of flagellated taxa to perform robust statistical comparisons. We annotated genomes (N = 21,551) into COG categories using eggNOG-mapper v2 [75], and calculated the genome size-corrected prevalence of each COG category per genome. We also investigated general genomic features such as genome size and the 16S rRNA gene copy number for each of the genomes to compare these genomic attributes between motile and non-motile taxa within each phylum. We included 16S rRNA gene copy number as it is considered a proxy for maximal potential growth rates in bacteria [45]. We compiled and identified the genes involved in chemotaxis (Supplementary Data 5) across the genomes of flagellated and non-flagellated taxa as a validation given the chemotaxis signaling pathway is an activator of the flagellar motor system [4].

### Estimation of the prevalence of flagellar motility in microbial communities using metagenomic information

We applied a method to estimate the prevalence of flagellar motility as a community-aggregated trait using metagenomic information from bacterial communities (Figure 3A). To this end, we first assembled ‘mock’ metagenomes containing different proportions of genomes from flagellated and non-flagellated taxa from the subset we originally used for boosted regression model training. We selected 20 genomes of taxa with empirically verified flagellar motility capabilities spanning the phyla Proteobacteria, Firmicutes, and Actinobacteria as these are ubiquitous taxa and are well-represented in our training data. We used ART (a next-generation sequencing simulator; [76]) to simulate short (150bp) shotgun sequencing reads at a coverage of 50% of these genome mixtures. We did not choose higher coverage as soil metagenomic datasets do not usually exceed 50% community coverage [77]. We then constructed a DIAMOND (v2.0.7; [78]) database containing the protein variants for each of the 21 selected genes (7-633 variants per gene) identified from the genomic analyses (see above) that were determined to be robust predictors of flagellar motility, as well as the variants contained in GTDB for the 120 single-copy marker genes that constitute the taxonomic basis of GTDB [27]. We annotated the simulated metagenomes using blastx (v2.13.0; [79]) on this custom protein database. In this way, we obtained a reads-per-kilobase (RPK) index for both the flagellar gene and the single-copy marker gene sets by taking the median gene length-corrected number of hits of each protein across the 21 and 120 unique proteins, respectively. We finally built a ‘flagellar motility index’ based on the ratio between the flagellar gene RPK and the single-copy marker gene RPK (see overview in Figure 3A). The use of single-copy marker genes in this manner offers a general normalization of the flagellar gene read count – which can vary due to differences in library size, coverage, or diversity – as these single copy genes are assumed to be present in every bacterial genome [80]. We then determined the linear relationship between the ‘flagellar motility index’ and the proportions of genomes that were able to produce flagella across mock metagenomes, following a similar approach to [48]. We used this standard curve to estimate the proportion of genomes in a given metagenome that are able to produce flagella based on the ‘flagellar motility index’, as expressed in equation *(1)*:

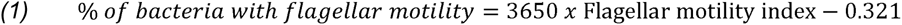

where the ‘flagellar motility index’ is the ratio between the median RPK of the 21 flagellar motility genes over the median RPK of the 120 single-copy marker genes (Figure 3A). This method allows the estimation of the prevalence of flagellar motility in any given bacterial metagenome based on the assumption that flagellar genes are usually found in single copies among bacterial genomes [23].

### Testing associations between bacterial flagellar motility and soil carbon availability

We selected metagenomic datasets that covered expected gradients in soil C availability, which we hypothesized to be positively associated with bacterial flagellar motility. Soil C availability is challenging to measure *in situ* and direct measurements of soil C availability (which is not equivalent to total C concentrations) are rarely compiled along with metagenomic data. We thus selected datasets that we expect based on published research to span gradients in C availability, recognizing that C availability is often correlated with other soil variables. The datasets included are the following: 1) soils from across the USA spanning gradients in soil depth (surface, 0-20; subsurface, 20-90cm, N = 66), where total organic C decreases with depth [51]; 2) a net primary productivity (NPP) gradient across Australia (N = 38, [81]), where higher NPP is expected to be associated with higher soil organic C availability [52]; 3) a global comparison of rhizosphere and bulk soils associated with citrus plants (N = 20, [55]), where we would expect C availability to be higher in rhizosphere soils than in bulk soils [82]; and 4) a pot experiment with controlled water inputs comparing the rhizosphere and adjacent bulk soil of wheat plants (N = 24, [56]).

Since all these datasets contain factors that likely covary with soil C availability, we additionally obtained metagenomic data from soils that were incubated with or without glucose amendments over a 117-d incubation period in a previous study [57]. In this experiment, glucose was added weekly to sub-samples of a single soil at a rate of 260 μg C g dry wt soil^−1^ day^−1^ (see [57] for full details). Since this experiment was performed under constant moisture conditions and in the absence of plants [57], the glucose amendments should lead to an increase in C availability with minimal direct effects on other soil attributes. The addition of glucose in this experiment led to a 7.9-fold increase in the microbial CO_2_ respiration rates [57], confirming that the C available to soil microbes increased in the soils amended with glucose compared to the controls (i.e. soils that received only an equivalent amount of water).

We then generated metagenomic data from the 9 soil samples harvested from the glucose amendment experiment. For each soil sample (4 with added glucose, 5 without glucose), we used 0.25g of soil for DNA extraction using the DNeasy PowerSoil Pro tube kit (Qiagen). The shotgun sequencing library was prepared using Illumina’s DNA Prep kit and Unique Dual Indexes (Illumina, CA). Samples were quantified using Qubit and pooled at equimolar concentrations. The library was run on a NovaSeq 6000 (Illumina, CA) at the Texas A&M AgriLife Genomics & Bioinformatics Service (USA) using a 2×150 cycle flow cell. Sequence cluster identification, quality prefiltering, base calling and uncertainty assessment were done in real time using Illumina’s NCS 1.0.2 and RFV 1.0.2 software (Illumina, CA) with default parameter settings. We also analyzed the 16S rRNA gene sequencing information on the same soil communities (see [57] for details on how this data was generated and processed).

### Processing of shotgun metagenomic sequencing reads from datasets covering gradients in soil C availability

To process the metagenomic data from all of the datasets described above (157 metagenomes in total), we first downloaded the sequences from the Sequence Read Archive (SRA) of NCBI when applicable, and ran trimmomatic (v0.39; [83]) to remove adapters and low quality base pairs using a phred score of 33 as a threshold, only keeping reads above 100bp after trimming. We used blastx on the custom DIAMOND database we created to annotate the metagenomic reads. We filtered out reads that had <50% bit score, <60% identity to the reference protein, and an e-value higher than 0.001. We finally measured the flagellar motility index based on the ratio between the median reads-per-kilobase (RPK) of the flagellar genes and the median RPK of the 120 single-copy marker genes as described above, and fitted equation *(1)* to quantify the prevalence of flagellar motility in any given metagenome using the method outlined in Figure 3A.

### Statistical analysis

All statistical analyses were conducted in R (v4.1.3; [84]). We used principal components analysis (PCA) to visualize how well the presence/absence of the selected flagellar motility genes was able to discriminate between the genomes of flagellated and non-flagellated taxa. To identify potential differences in the life history strategies of flagellated and non-flagellated taxa, we used multiple Mann-Whitney U tests with Bonferroni correction for multiple comparisons to investigate whether particular COG categories were overrepresented in genomes from flagellated versus non-flagellated taxa. The results were presented as the log2-fold ratio. We used Mann-Whitney U tests to investigate associations between flagellar motility and the 16S rRNA gene copy number and the number of chemotaxis genes in any given genome due to non-normality of the data. We compared differences in genome size between flagellated and non-flagellated taxa using Welch two-sample t-tests.

To test for differences in the prevalence of flagellar motility between surface and subsurface soils we used a mixed effects linear model with location coded as random factor, and for the test between rhizosphere and bulk soils in wheat we used Welch two-sample t-tests. Since we only had a single rhizosphere and bulk soil observation per site, in the global citrus rhizosphere dataset we first calculated the difference in the prevalence of flagellar motility in the rhizosphere over bulk soil at each site, and then tested whether these differences were significantly different from zero using a one-sample t-test. These tests were implemented using different arguments of the *t.test* function in base R [84]. We used Pearsons’ correlations to evaluate relationships between the prevalence of flagellar motility and NPP, and used linear regression to represent the standard curve to quantify the prevalence of flagellar motility in metagenomes.

Finally, we used 16S rRNA gene sequencing information from samples in the glucose amendment experiment [57] to investigate the shifts in the taxonomic composition of soil bacterial communities upon glucose addition. Specifically, we investigated which bacterial Amplicon Sequence Variants (ASVs) responded to glucose addition using ANCOM-BC [85]. The taxonomic composition of these bacterial communities was investigated using the phyloseq R package (v1.38.0; [86]), and we tested the effect of glucose amendment on the prevalence of flagellar motility assessed using our metagenome-based method using the Welch two-sample t-test.

## Acknowledgements

We thank Michael Hoffert and Thomas B. N. Jensen for assistance with the bioinformatical analyses, and Jessica Henley, Caihong Vanderburgh, and Jordan Galletta for generating the metagenomic sequence data.

## Author contributions

JR and NF conceived and designed the study. JR performed the data analyses. KF, JML, HC, AB, and MSS contributed data to the study. JR and NF wrote the manuscript, with input from all co-authors.

## Supplementary Material

**Supplementary Figure 1.**
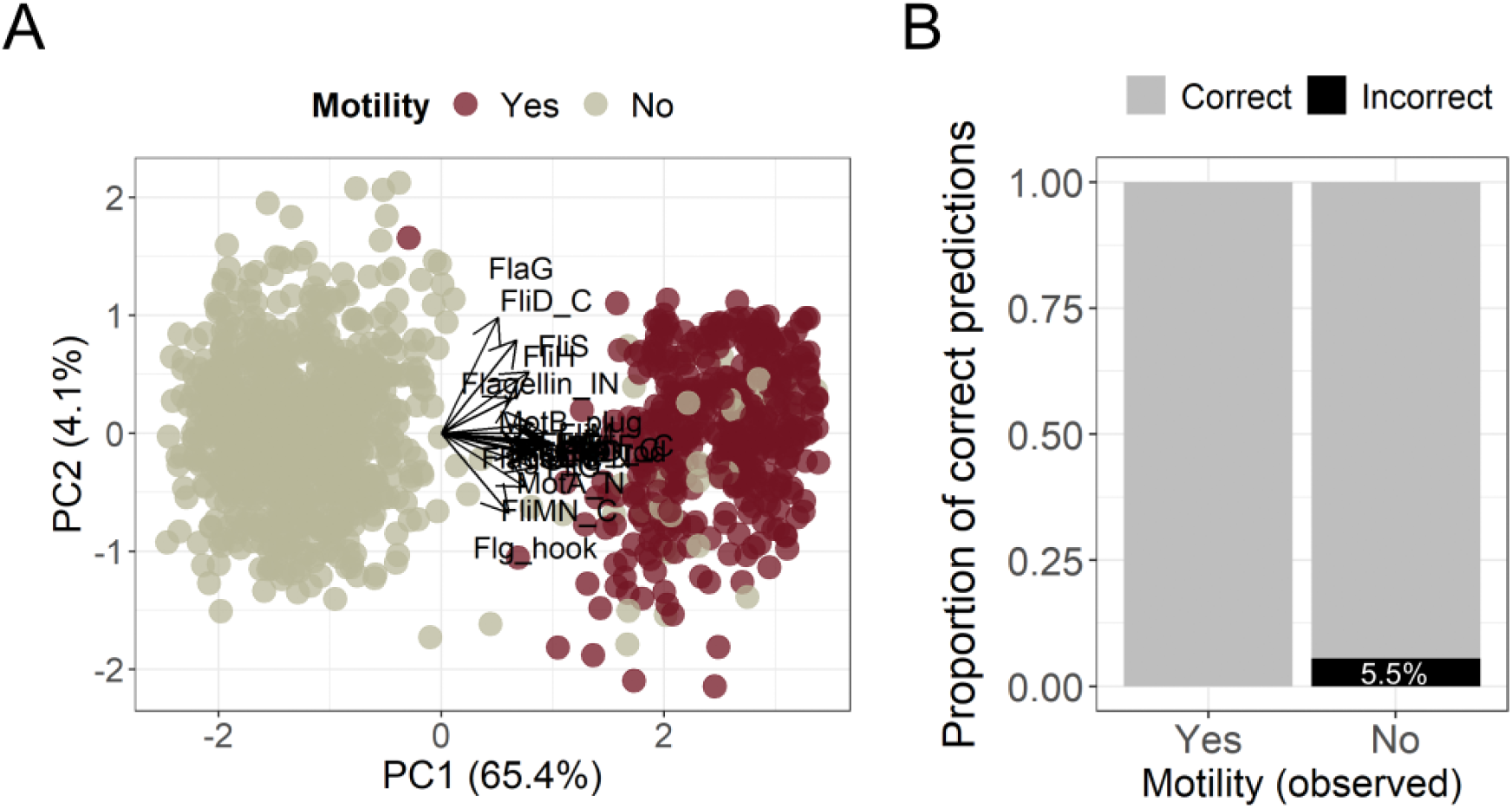
Prediction of the capacity to display flagellar motility in bacterial taxa based on presence/absence information of 21 genes involved in flagellar assembly. A. Principal Components Analysis (PCA) based on the presence/absence of 21 flagellar genes in genomes of taxa that have been empirically found to be flagellated (N = 388 genomes) or non-flagellated (N = 837 genomes). Empirical information on flagellar motility was obtained from the bacterial phenotypic trait data compiled in [26]. We only included genomes that were 100% complete, contained an assembled 16S rRNA gene, and showed no signs of chimerism. B. Accuracy of a boosted regression machine learning model trained on the genomes shown in panel A for the prediction of the capacity for flagellar motility in any given bacterial genome based on the presence/absence of 21 flagellar genes. Accuracy was tested on 30% of the original genome set (116 genomes from flagellated taxa and 251 genomes from non-flagellated taxa). Genomes were obtained from the Genome Taxonomy Database (GTDB r207; [27]).

**Supplementary Figure 2.**
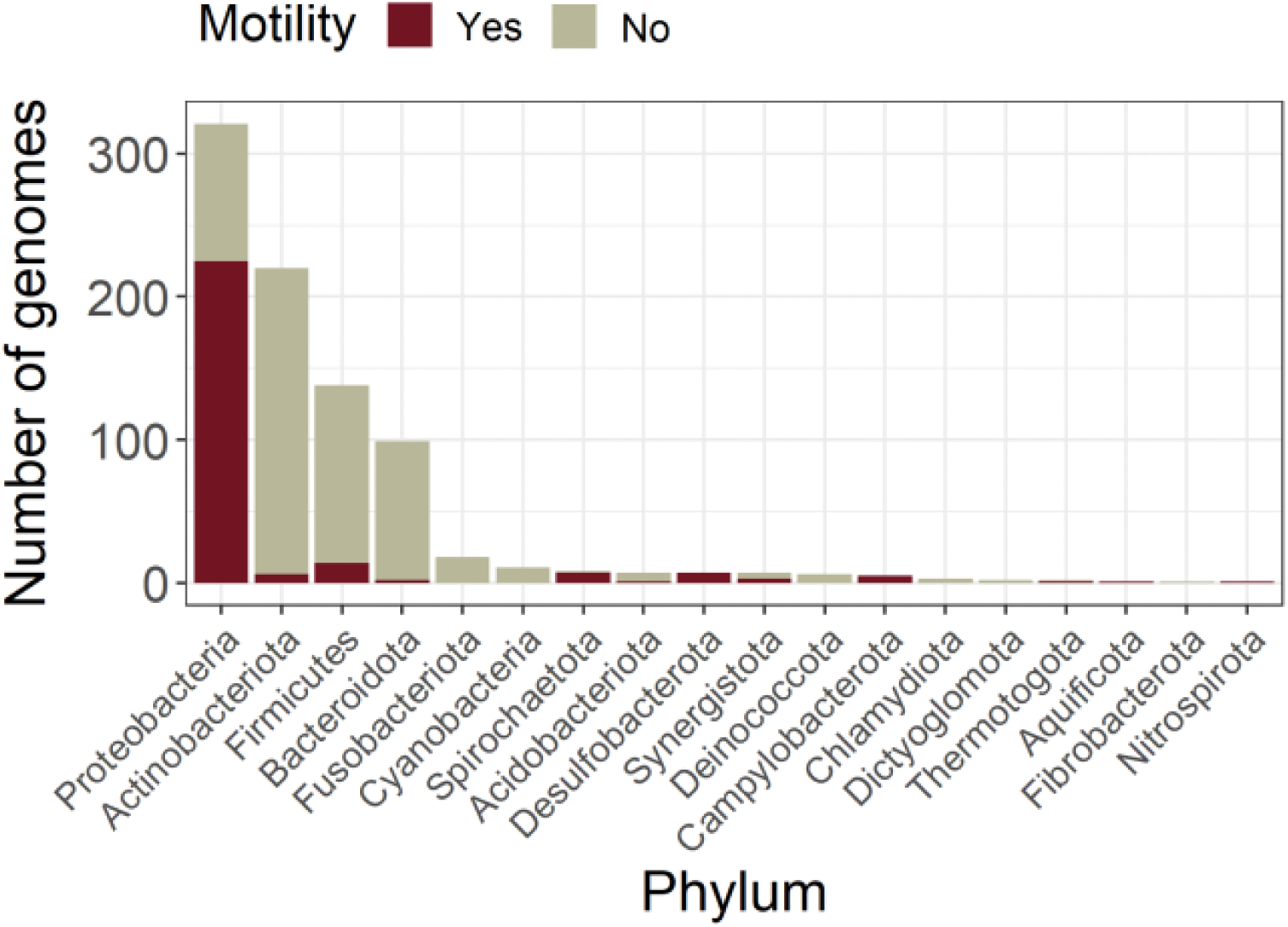
Taxonomic distribution of genomes with empirically determined capacity for flagellar motility that were used as training data for a boosted regression machine learning model to predict the capacity for flagellar motility based on the presence/absence of 21 flagellar genes. Flagellar motility information was obtained from the bacterial phenotypic trait data compiled in [26]. N_Training set_ = 858 genomes, N_Full set_ = 1225.

**Supplementary Figure 3.**
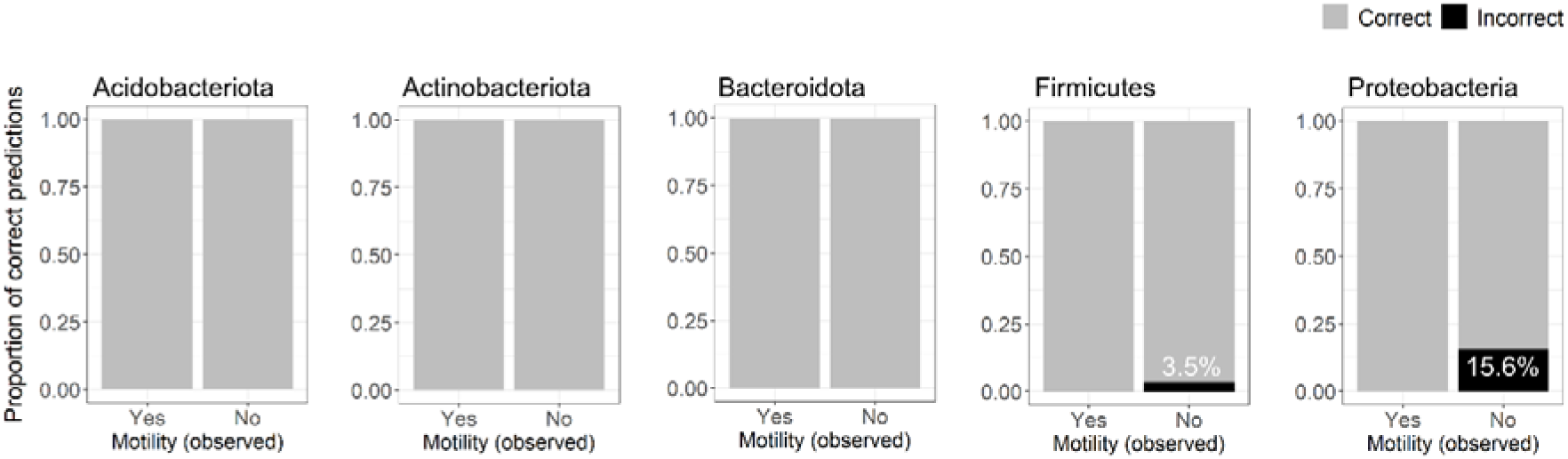
Predictive accuracy across phyla of a boosted regression machine learning model for the prediction of the capacity for flagellar motility in any given bacterial genome based on the presence/absence of 21 flagellar genes. Accuracy was tested on 30% of the original genome set (116 genomes from flagellated taxa and 251 genomes from non-flagellated taxa). Genomes were obtained from the Genome Taxonomy Database (GTDB r207; [27]). N_Acidobacteriota_ = 8, N_Actinobacteriota_ = 87, N_Bacteroidota_ = 52, N_Firmicutes_ = 66, N_Proteobacteria_ = 123.

**Supplementary Figure 4.**
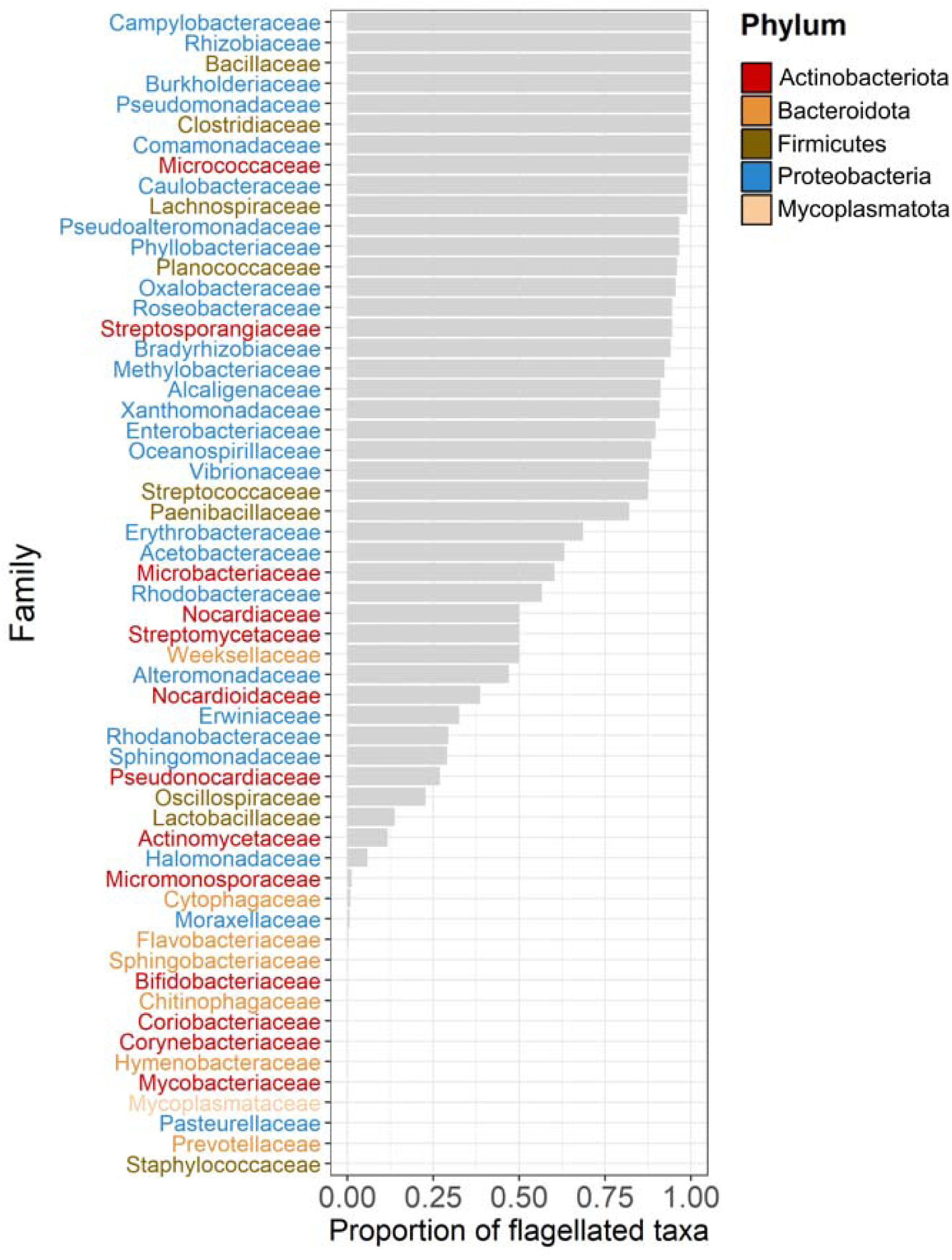
Prevalence of flagellar motility across bacterial families containing more than 100 high-quality genomes in the Genome Taxonomy Database (GTDB r207; [27]). We only included genomes that were >95% complete, contained an assembled 16S rRNA gene, and showed no signs of chimerism. N = 23,256 genomes.

**Supplementary Figure 5.**
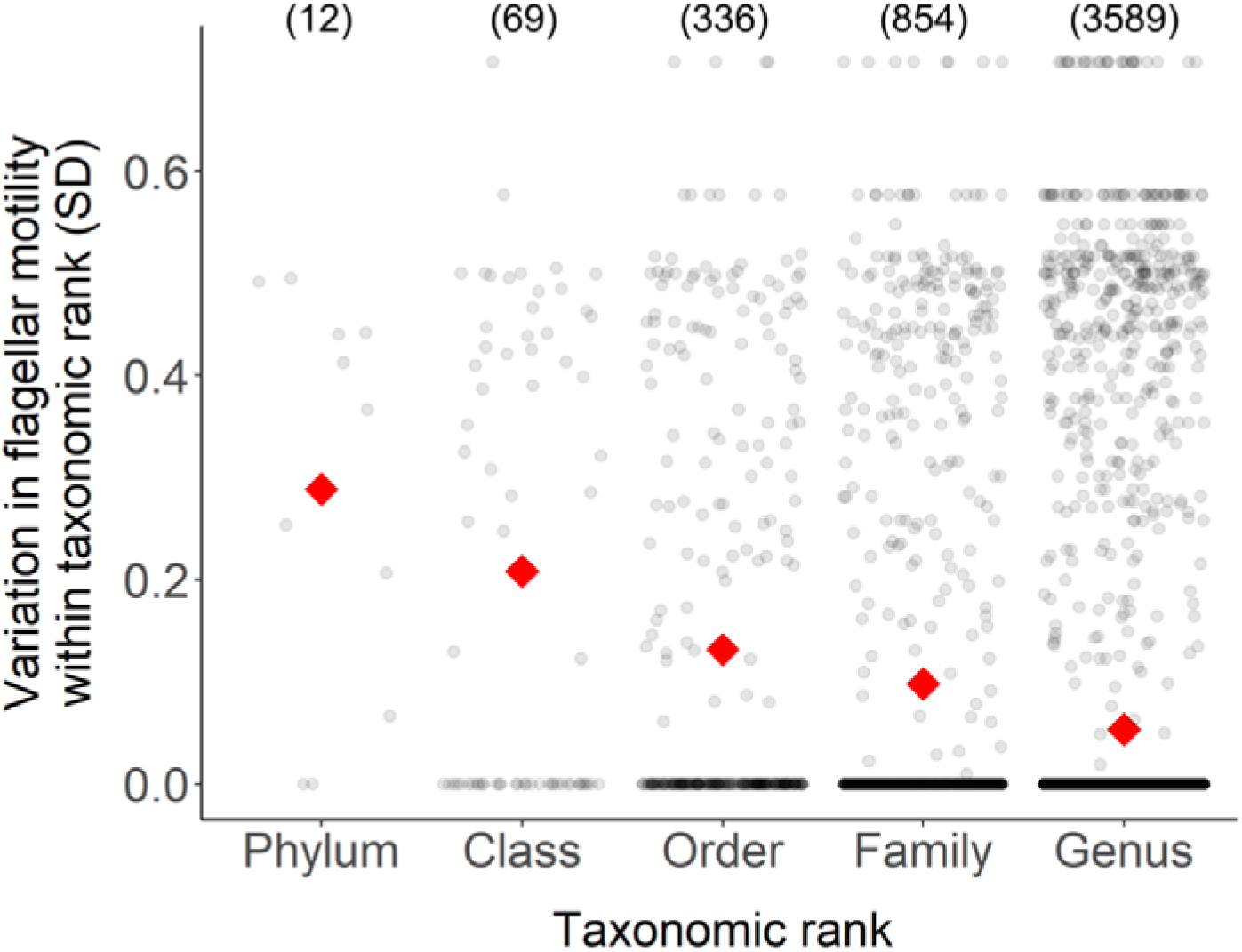
Variation in flagellar motility status across taxonomic ranks. A measure of variation in the flagellar motility status of taxa belonging to different taxonomic ranks was obtained from the standard deviation (SD) of their flagellar motility status (1, flagellated; 0, non-flagellated). Numbers in brackets indicate the total number of unique taxa within each of the taxonomic ranks. Red diamonds indicate the mean of the standard deviation of the flagellar motility status within each taxonomic rank.

**Supplementary Figure 6.**
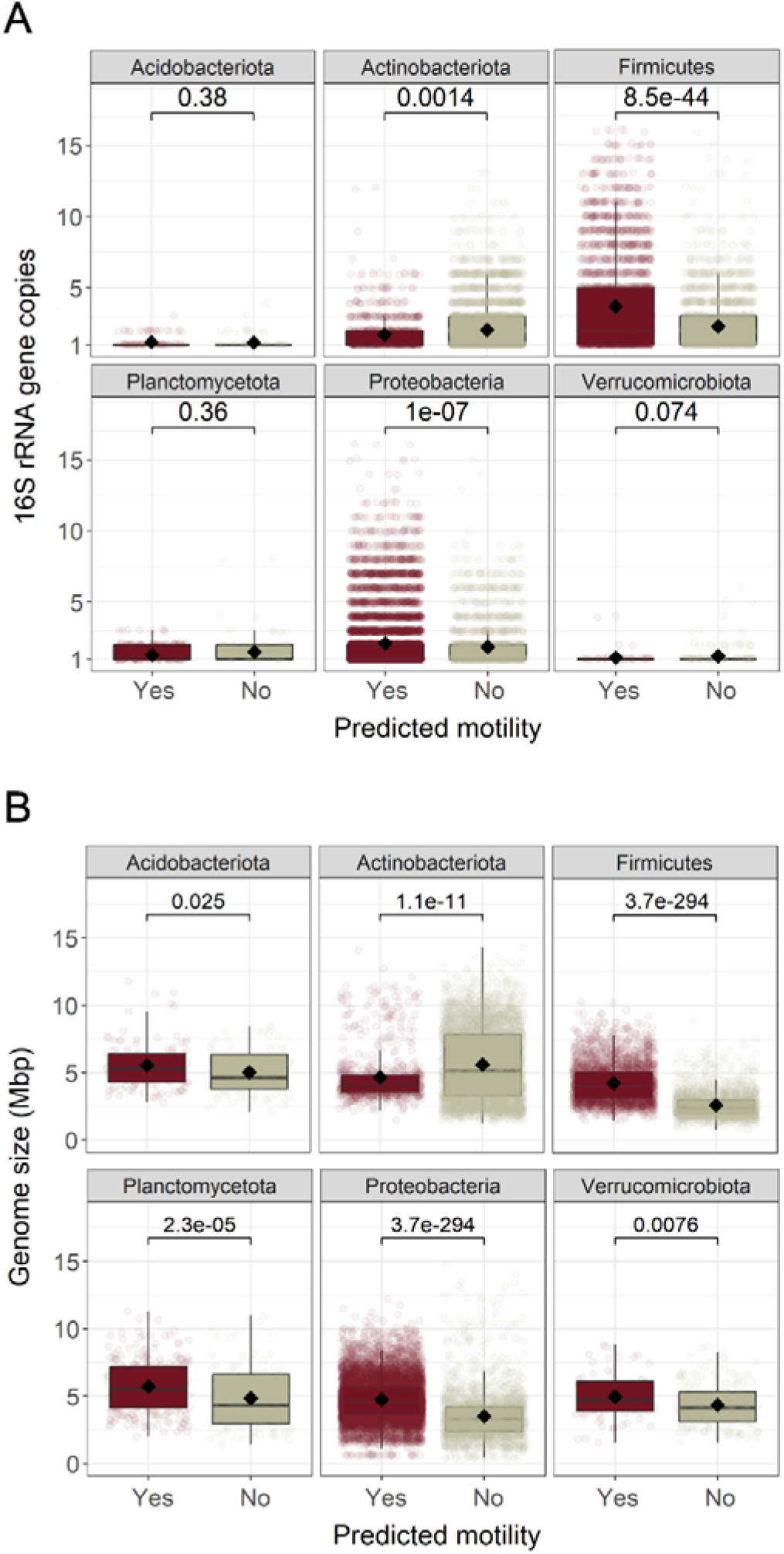
Distribution of the total number of 16S rRNA gene copies per genome and genome size across the 6 phyla with even proportions of taxa predicted to be flagellated and non-flagellated. A. Number of 16S rRNA gene copies in genomes of taxa predicted to be flagellated and non-flagellated. B. Genome size of taxa predicted to be flagellated and non-flagellated. Statistical significance was obtained from Mann-Whitney U tests (P < 0.05), N = 21,551.

**Supplementary Figure 7.**
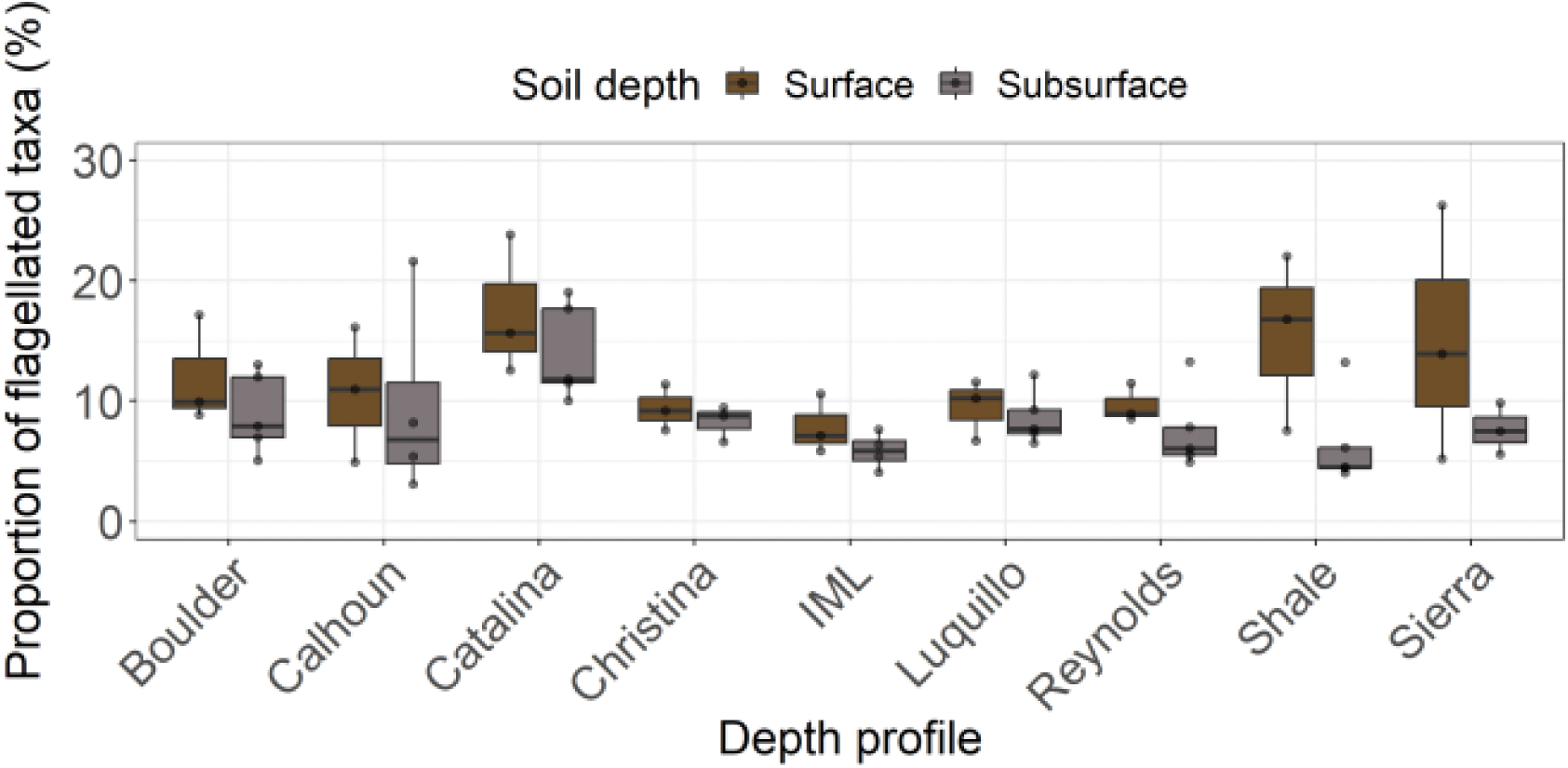
Estimated prevalence of flagellar motility in bacterial communities from 9 soil depth profiles collected across the USA (Surface, 0-20cm; Subsurface, 20-90cm, N = 66; [51]).

**Supplementary Figure 8.**
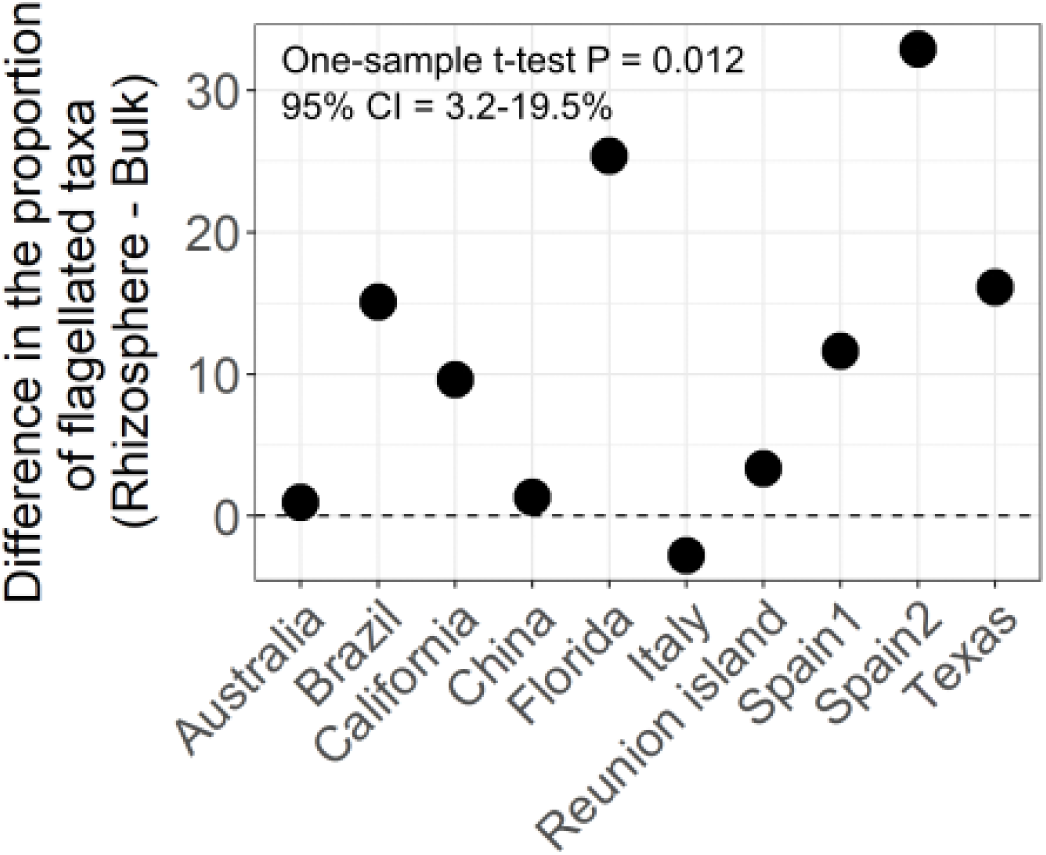
Difference in the prevalence of flagellar motility in rhizosphere and bulk soil bacterial communities collected from citrus species across the globe (N = 10; [55]).

**Supplementary Figure 9.**
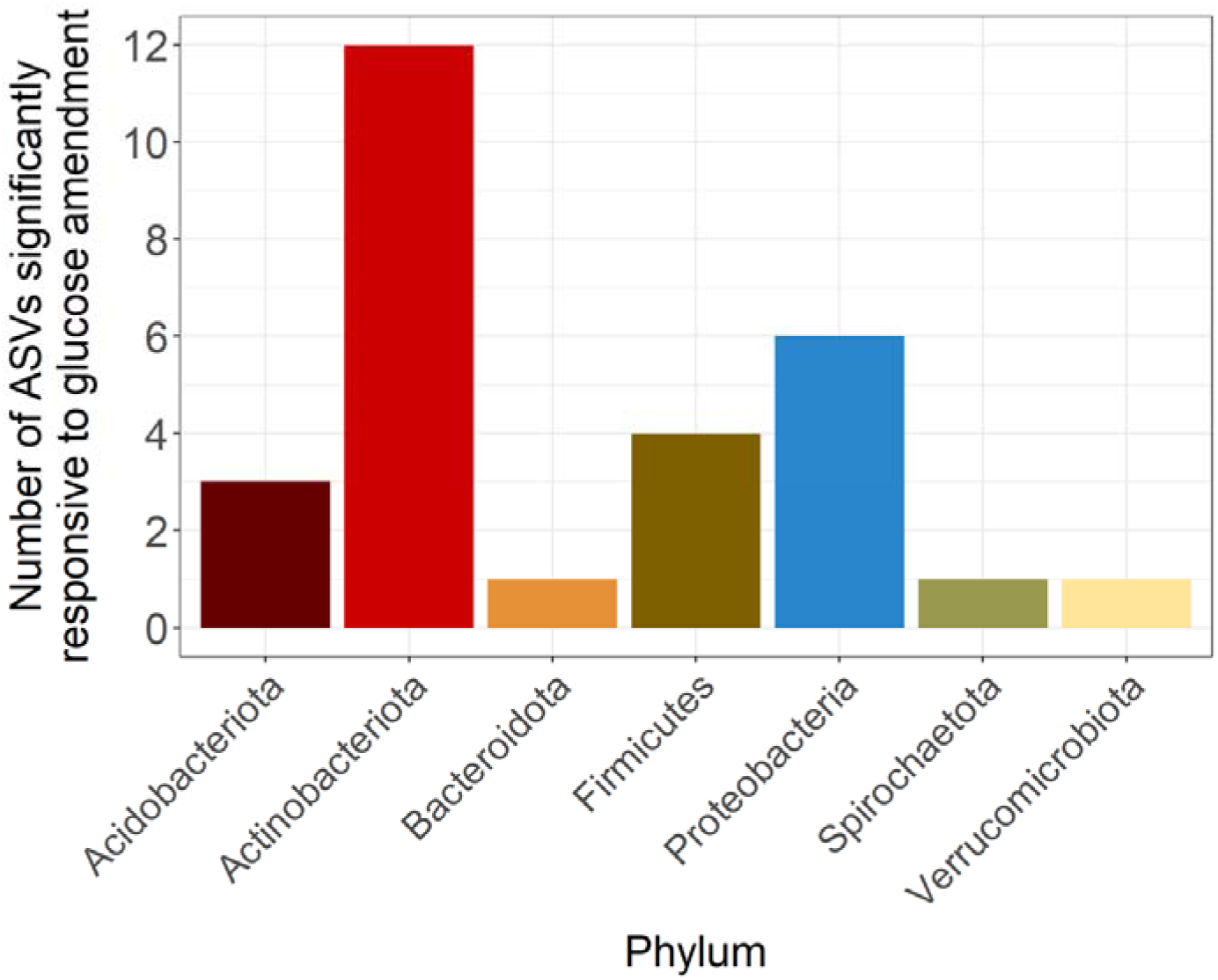
Taxonomic composition of the Amplicon Sequence Variants (ASVs) that responded to glucose amendment in soil. Bacterial communities from a 117 day soil incubation experiment with daily glucose amendment were characterized using amplicon sequencing of the 16S rRNA gene [57]. ASVs were considered responsive to glucose amendment based on a differential abundance analysis comparing bacterial communities from soils amended with glucose versus communities from soils that did not receive any external carbon inputs (N = 28 responsive ASVs).

